# Controlling False Discovery in CRISPR Screens

**DOI:** 10.1101/2025.04.24.650434

**Authors:** Joshua Dempster, Barbara De Kegel, Alex Kalinka, Francisca Vazquez, Catarina D Campbell

**Affiliations:** The Broad Institute, 451 Main St, Cambridge, MA; Cancer Research Horizons, Joint AstraZeneca-Cancer Research Horizons Functional Genomics Centre, Cambridge, UK

**Keywords:** CRISPR screen, false discovery, anchor screen, cancer cell line, genome editing

## Abstract

Excluding false positives is critical for interpreting CRISPR screens. Here, we introduce a new Chronos module for estimating false discovery rates for identifying knockouts that cause loss of viability or have differential viability effects in different conditions. We introduce a rigorous benchmarking framework using real CRISPR data. We show with multiple real datasets that existing methods such as MAGeCK are miscalibrated and can generate uncontrolled numbers of false positives even after multiple hypothesis correction. Only Chronos correctly controls false discovery for all tested tasks. Additionally, Chronos’s estimates are well-calibrated, allowing users to accurately specify the acceptable false discovery rate.

## Background

Lack of reproducibility is a widespread problem in preclinical research and contributes to the inefficiency of drug discovery efforts(1). There are many reasons why an experiment may fail to reproduce; one of the simplest and most addressable is a statistical failure to control false discovery in the analysis of large-scale datasets(2). Improper control of false discovery can be due to: comparison to a rigid or incomplete null hypothesis, which may assume errors are normal when they are kurtotic or independent when they covary; overlooking sources of bias; ignoring the multiple hypotheses that were implicitly tested through researcher degrees of freedom; or failure to imagine a critical mode of error in the data.

Large- scale pooled CRISPR screens have been widely adopted for interrogating gene function(3) and have become common sources of nomination for therapeutic targets, particularly in oncology. Compared to previous genetic perturbation methods, CRISPR shows much greater specificity and potency than RNA interference (RNAi)(4), and more potency and flexibility than transcription activator-like effector nucleases (TALENs)(5). However, as CRISPR screens have proliferated, biologists have developed a greater appreciation for the complexities and possible artifacts of the data. Tools for estimating or correcting gene viability effects from CRISPR data include MAGeCK(6), MAGeCK-MLE(7), CERES(8), JACKS(9), CRISPRCleaner(10), BAGEL(11), BAGEL2(12), CRISPRBetaBinomial(13), DrugZ(14), and Chronos(15). Some of these methods return *p*-value estimates, but to our knowledge these have not been independently benchmarked for their ability to control false discovery.

In this work, we present methods for controlling false discovery in CRISPR screens that incorporate multiple sources of error, known and unknown, in the null hypothesis. CRISPR screens are frequently performed to answer one of two questions: 1. which gene losses cause true change in the viability phenotype; and 2. which gene losses have different viability impacts in different conditions. These questions are implicitly statistical, as they require making some determination of what constitutes a true discovery among noisy data. Of the tools mentioned above, MAGeCK, JACKS, DrugZ and CRISPRBetaBinomial offer intrinsic methods for estimating the significance of a change in viability. These methods assess whether the readcounts of an sgRNA in a condition differ significantly from a control condition; the control condition can be an early timepoint or plasmid abundance to answer question 1, or e.g. an unperturbed condition to answer question 2. However, we will show below that these existing methods can be dramatically miscalibrated and therefore fail to control false discovery. Here, we introduce a new suite of methods in Chronos that control false discovery in the tested settings while better isolating true findings.

## Results

Determining true loss of viability within a condition and determining significant differences in viability between conditions are different problems. Comparing conditions requires comparing two screens. In addition to being intrinsically noisier, the difference between screens is affected by biases in cell representation, growth rate, and knockout efficacy that do not apply to measuring viability within a single screen. However, existing methods other than DrugZ use the same statistical methodology for both of these tasks. To address the first task, we present two methods, frequentist and empirical-Bayesian, either of which may be more appropriate depending on the nature of the experiment (see Discussion). Both methods accept a matrix of gene effect estimates, such as that produced by the methods above, to produce an estimated false discovery rate for the hypothesis that a gene knockout causes loss of viability *vs* the null hypothesis that it does not within each cell line.

The frequentist method uses negative (nonessential) control genes to create an empirical null distribution of effects (**Fig. 1a**). For a gene of interest, we compute how many negative controls have more negative gene effects than the gene of interest, from which it is straightforward to compute an empirical *p-*value(16) (**Fig. 1b**). We estimate the false discovery rate (FDR) using the two-stage Benjamini-Hochberg algorithm(17). The power of the frequentist method is that it implicitly captures all the forms of bias or noise, such as copy number effect or off-target effects, whether or not the experimenter knows of them. However, the smallest possible *p*-value that can be produced with this method is 1 / (the number of negative controls). In a genome-wide screen, the set of unexpressed genes provides thousands of high-confidence negative controls. But in a subgenome library with a small number of negative controls, this limit may prevent any genes from reaching significance.

**Figure 1:**
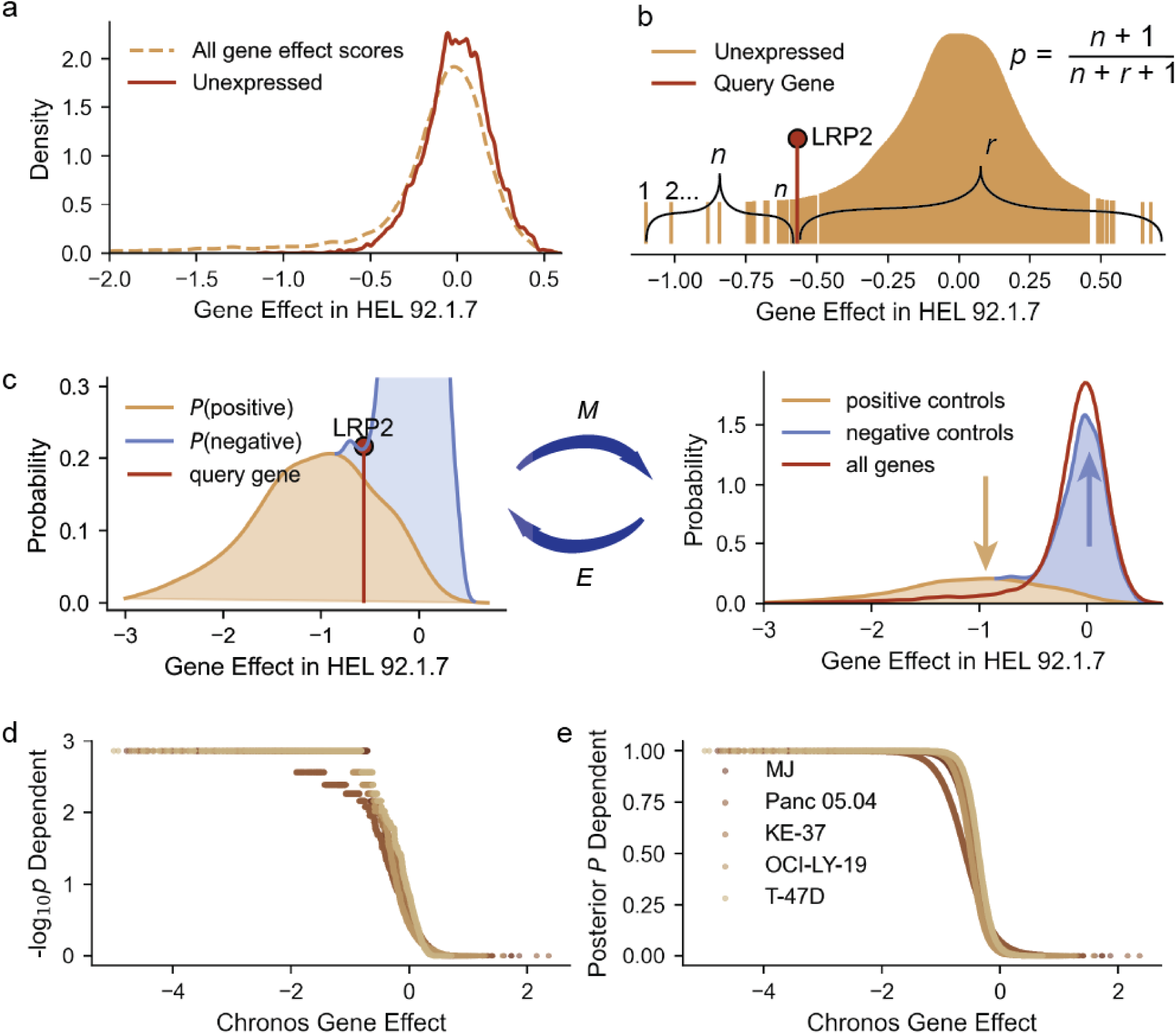
Calling significant depletion. **a.** The distribution of all gene effect scores and unexpressed gene effect scores in the cell line HEL 92.1.7. **b.** Schematic for the empirical *p*-value for the gene LRP2 using unexpressed genes as the null distribution, where *n* is the number of negative controls with more negative effects than the query gene and *r* the number with more positive effects. **c**. Illustration of the Bayesian posterior probability estimation. In the *M* step (left), the posterior probability of each gene being generated from the positive control distribution is updated, while in *E* (right) the estimate of the fraction of all gene scores drawn from the positive vs the negative control distribution is updated. **d.** Relationship of empirical *p*-values to gene effect scores for five lines screened in DepMap with the Avana library. **e.** Relationship of posterior probabilities to gene effect scores.

The empirical Bayesian method is complementary to the frequentist. In a specified cell line, given a set of positive (essential) control genes and negative control genes, we first estimate the probability density function (PDF) of each group using a Gaussian kernel. This step is similar to the first step of BAGEL(11,12). We take the minimally informative prior that each gene has equal probability of being generated from the positive or negative control distributions. We then estimate posterior probabilities using the PDFs and Bayes’ rule (*E* step), followed by updating the estimated total proportions of the data belonging to the positive and negative control distributions (*M* step), and repeat until convergence (**Fig. 1c**). The posterior probabilities can be converted to a positive false discovery estimate by arranging them in decreasing order and taking one minus the cumulative mean(18). Because the control samples are finite, density estimates in the tails can become quite noisy; therefore we regress the probability of dependency on the observed gene effect using a sigmoid function (see Methods). We chose five random Avana screens conducted by the Cancer Dependency Map to illustrate the relationships of *p*-value and posterior probability to gene effect **(Fig. 1de**).

Most tools for inferring gene effects from CRISPR data do not include methods to estimate significance. Thus, for comparison, we selected available tools that estimate significance using different approaches: MAGeCK, CRISPRBetaBinomial, and JACKS. The tested methods are summarized in Table 1. The updated MAGeCK-MLE algorithm includes two methods for computing *p*-values: the Wald method, which assumes that the *β* (gene effect) parameter has *χ*^2^-distributed errors(7); and the permutation method, which shuffles the assignment of sgRNAs to genes. The CRISPRBetaBinomial (CB2) method assumes that CRISPR readcounts are drawn from the beta-binomial distribution(13). Finally, JACKS estimates guide efficacy assuming normality of log-fold change in sgRNA abundance. JACKS computes *p*-values by accepting a list of negative control genes and shuffling sgRNA assignment to the negative controls. Since only some of these methods include corrections for copy-number toxicity(19), we did not use any copy number correction.

**Table 1:** Summary of methods. Task 1: identify gene knockouts causing significant depletion within a screen. Task 2: identify genes with significantly different viability effects in two different conditions. H0: null hypothesis. i.i.d.: independently and identically distributed. H1: alternative hypothesis.

| Code | Method | Task | Statistical Assumptions/Requirements |
| --- | --- | --- | --- |
| MW | MAGeCK-MLE<br>Wald | 1 and 2 | Normality of inferred $\beta$ parameter under H0 |
| MP | MAGeCK<br>permutation | 1 and 2 | sgRNA readcounts for each gene are i.i.d. |
| CB2 | CRISPR<br>beta-binomial | 1 and 2 | sgRNA log fold-changes are beta-binomially distributed under H0 |
| JK | JACKS | 1 and 2 | sgRNA log fold-changes are i.i.d. |
| DZ | DrugZ | 2 only | sgRNA log fold-changes are normal and i.i.d. after normalizing by variance of sgRNAs with similar counts |
| ChF | Chronos<br>frequentist | 1 only | Many known negative control genes that sample H0 |
| ChB | Chronos<br>Bayesian | 1 only | Known negative control genes that sample H0 and positive control genes that sample H1 |
| ChL | Chronos<br>likelihood | 2 only | Two independent biological replicates per condition |

We created training and testing sets of positive and negative control genes that were both mutually exclusive and nominated by orthogonal criteria (see Methods). We first evaluated control of false discovery by each method in a setting with no true discoveries. We ran the methods on the previously described five Avana screens while including only test and train negative control genes. For this comparison, the Chronos Bayesian method was excluded as it requires positive controls to be used. Methods accepting negative controls were supplied the training set, while all methods were evaluated on the test set. Of the methods, only Chronos exhibited good calibration, with *p*-values closely aligned to the expected uniform distribution (**Fig. 2ab**). All prior methods showed a marked excess of small *p-*values. Consequently Chronos made no false discoveries in these data, while the other methods made between 23 and 195 false discoveries (**Fig. 2c**).

**Figure 2:**
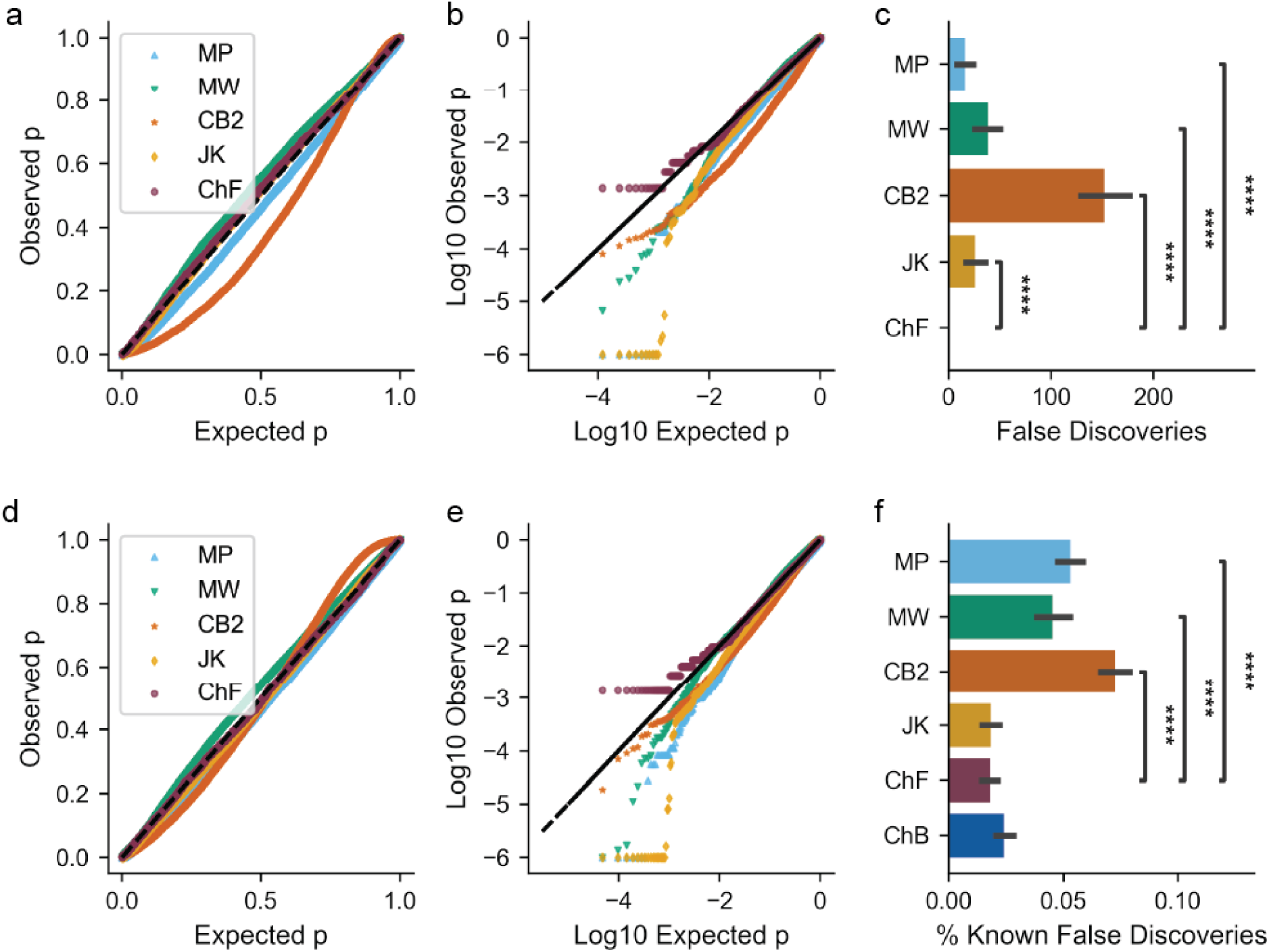
Calibration of false discovery control with different methods. **ab.** In a run with that only included 2,396 negative control genes for five chosen Avana screens, the observed distribution of *p* values vs the expected uniform distribution. MP: MAGeCK permutation, MW: MAGeCK Wald, CB2: CB2, JK: JACKS, ChF: Chronos frequentist. The Chronos Bayesian method requires positive controls and is excluded here. For visualization, *p*-values < 10^-6^ have been clipped to 10^-6^ **c**. Number of discoveries made in the test set of negative controls at estimated false discovery rate/positive false discovery rate (FDR) < 0.1 (all discoveries are false in this set). Error bars show confidence intervals using resampling with replacement. The Chronos Bayesian method does not produce *p*-values and is excluded here. The number of asterisks indicate the significance to that power of 10 using a *t-*test on bootstrap resampling. **de.** As above, but for a run that included all genes. *p*-values are included only for unexpressed genes. **f**: Fraction of total discoveries at FDR < 0.1 which are unexpressed. ChB: Chronos Bayesian.

Interestingly, JACKS has the worst calibrated *p*-values, but reports the lowest proportion of false discoveries of tested methods. This result arises from the definition of a false discovery rate used in JACKS. Rather than the traditional definition of expected proportion of true null hypothesis among all hypotheses of equal or more extreme statistics (20), an examination of its code reveals that JACKS defines the FDR at a given gene effect value as the ratio of the probability density of the negative controls to the probability density of all gene effects at that value. No estimate of the proportion of negative results is made, so both densities have unit area. This definition creates counterintuitive effects: JACKS’ FDR estimates are not necessarily monotonically related to its *p*-values, and its FDRs can and do exceed 1. Under certain conditions, notably that the amount of data goes to infinity while the number of true discoveries goes to zero and no gene knockouts *increase* viability, this definition will approximate the conventional definition of false discovery; but we have not found that those conditions hold in typical CRISPR screens (**Suppl. Fig. 1**).

We then evaluated control of false discovery in the same screens, now retaining all genes, and observed similar results (**Fig. 2de**). In this case, since there are real discoveries to be made, we expect even a perfectly calibrated method to report some false discoveries at a finite FDR. We find that both the frequentist and Bayesian Chronos methods report a similar fraction of unexpressed genes among their discoveries in this dataset (1.8%-2.4%) which is lower than MAGeCK or CB2 (4.5%-7.2%; **Fig. 2f**). JACKS again exhibits the worst *p*-value calibration but the lowest false discoveries of prior methods.

**Supplementary Figure 1:**
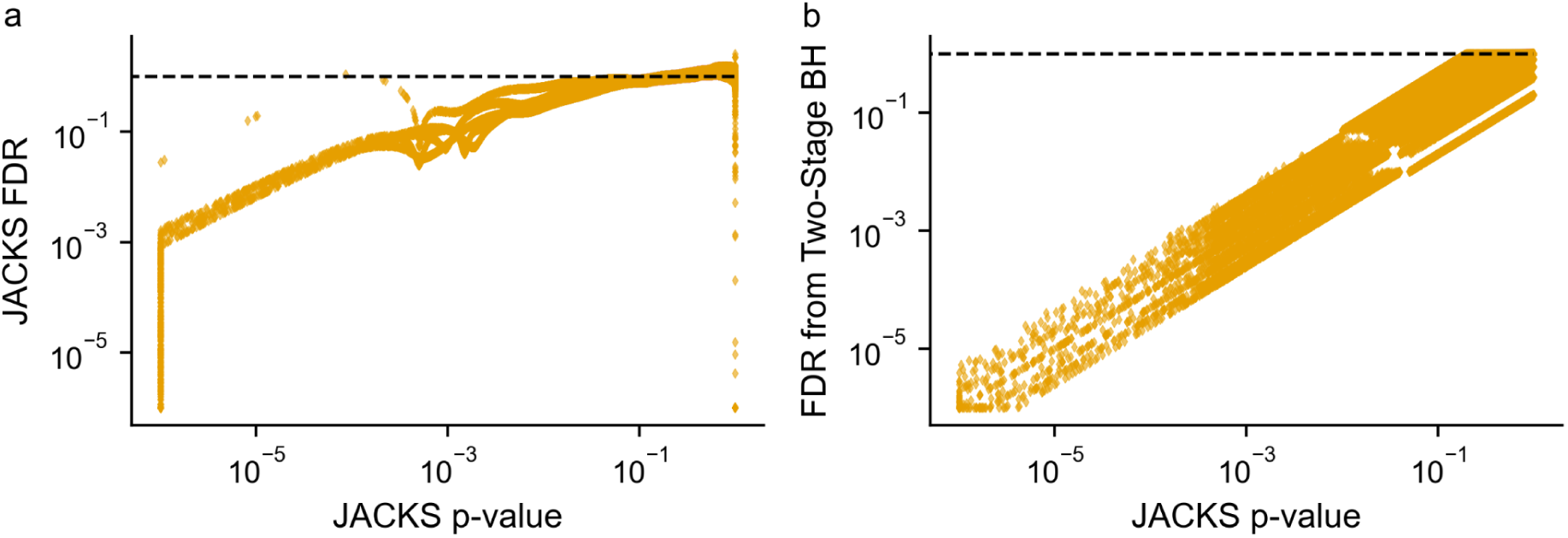
JACKS’ vs conventional FDRs. **a.** The relationship of JACKS’ FDRs to its *p*-values over the five chosen Achilles screens with all genes. The black line is drawn at FDR = 1. **b.** JACKS’ *p*-values vs a conventional two-stage Benjamini-Hochberg correction performed per cell line.

When examining how each method’s false discovery rate discriminates true from false discoveries in this data, we find both Chronos methods have very similar performance as evaluated by the area under the precision-recall curve (PR AUC) (0.65-0.66, vs 0.38-0.57 for competitors; **Fig. 3a**). Given that both Chronos methods rely on the presence of controls, we examined how discovery power is affected by decreasing numbers of controls. As expected, we find that the frequentist method loses power rapidly with less than 500 controls (**Fig. 3b**). When fewer than 100 negative controls are included, in most cases no positive controls could be identified at FDR < 0.1. In contrast, the empirical-Bayesian method maintains its power with as few as 10 positive and negative controls supplied. This led us to ask how well-calibrated the false discovery estimates of each method were; that is, how closely they matched the true false discovery rate. First, we subset the same screens to only positive (1,617) and negative (2,396) control genes before training. The Chronos Bayesian FDR estimates exhibited the least deviation from the true FDR (median absolute deviation: 0.013), followed by the Chronos frequentist FDRs (0.081) (**Fig 3c**). JACKS exhibits catastrophic miscalibration, which is unsurprising given its unusual definition of FDR.

**Figure 3:**
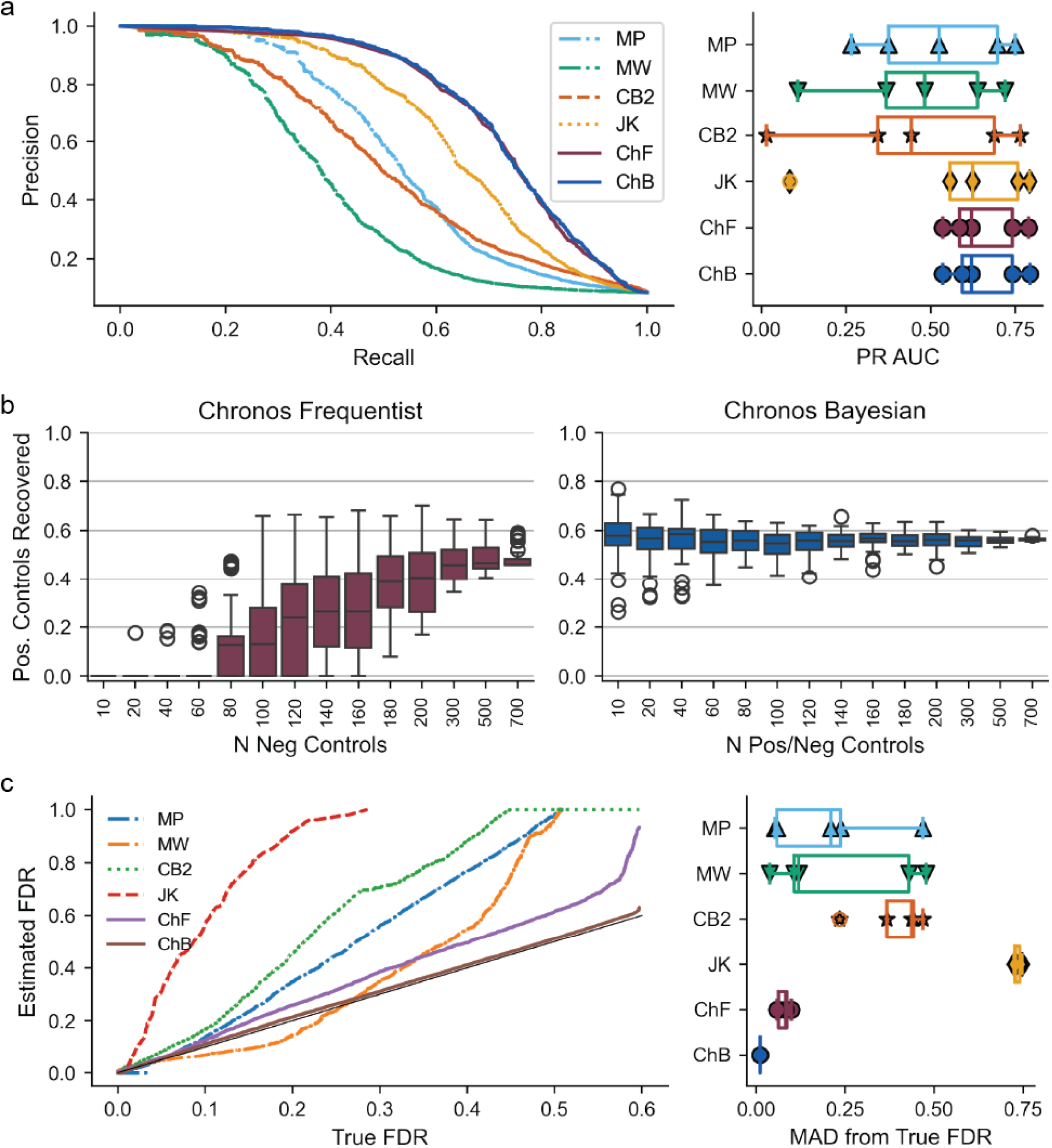
Control separation and calibration. **a.** Recall and precision for identifying expected common essential dependencies vs unexpressed genes using *p*-values (MP, MW, CB2, ChF) or posterior probabilities (ChB). Right: total area under the precision-recall curve for each cell line using each method. **b.** Downsampling analysis where *p* values are estimated 50 times using random non-replacement samples of the training negative controls (left) and posterior probabilities from 50 non-replacement samples of positive and negative controls (right), where the number of controls included is given on the *x*-axis. For each sample, recovery of test positive controls at FDR < 0.1 is reported on *y*. **c**. In a run with only positive and negative control genes, estimated FDR vs actual FDR. Estimated FDR values > 1 are excluded (JACKS). Right: median absolute deviation of the estimated FDR from the actual FDR for each cell line.

These results demonstrate that Chronos controls false discovery of dependencies, and further that it has the best discrimination between true and false discoveries. However, researchers often wish to understand how the viability effect of gene knockouts changes between two conditions (task 2). This question is more difficult, because our usual negative control groups such as unexpressed genes do not represent the full spectrum of gene dependencies and may fail to control false discovery for other types of genes, such as common essentials.

To get around this problem, we train multiple Chronos models (**Fig 4a**). In one model, we do not distinguish between replicates of the same cell line in different conditions. In other words, we generate one set of gene effect estimates for each cell line that combines both conditions. We call this the undistinguished model. We then train a model where we treat the combination of condition and cell line as a unique cell line, which we call the distinguished model. Our statistic is the increase in log likelihood of the observed readcounts with the higher-parameter distinguished model vs the undistinguished model. This statistic is reported individually for each cell line that is present in both conditions.

**Figure 4:**
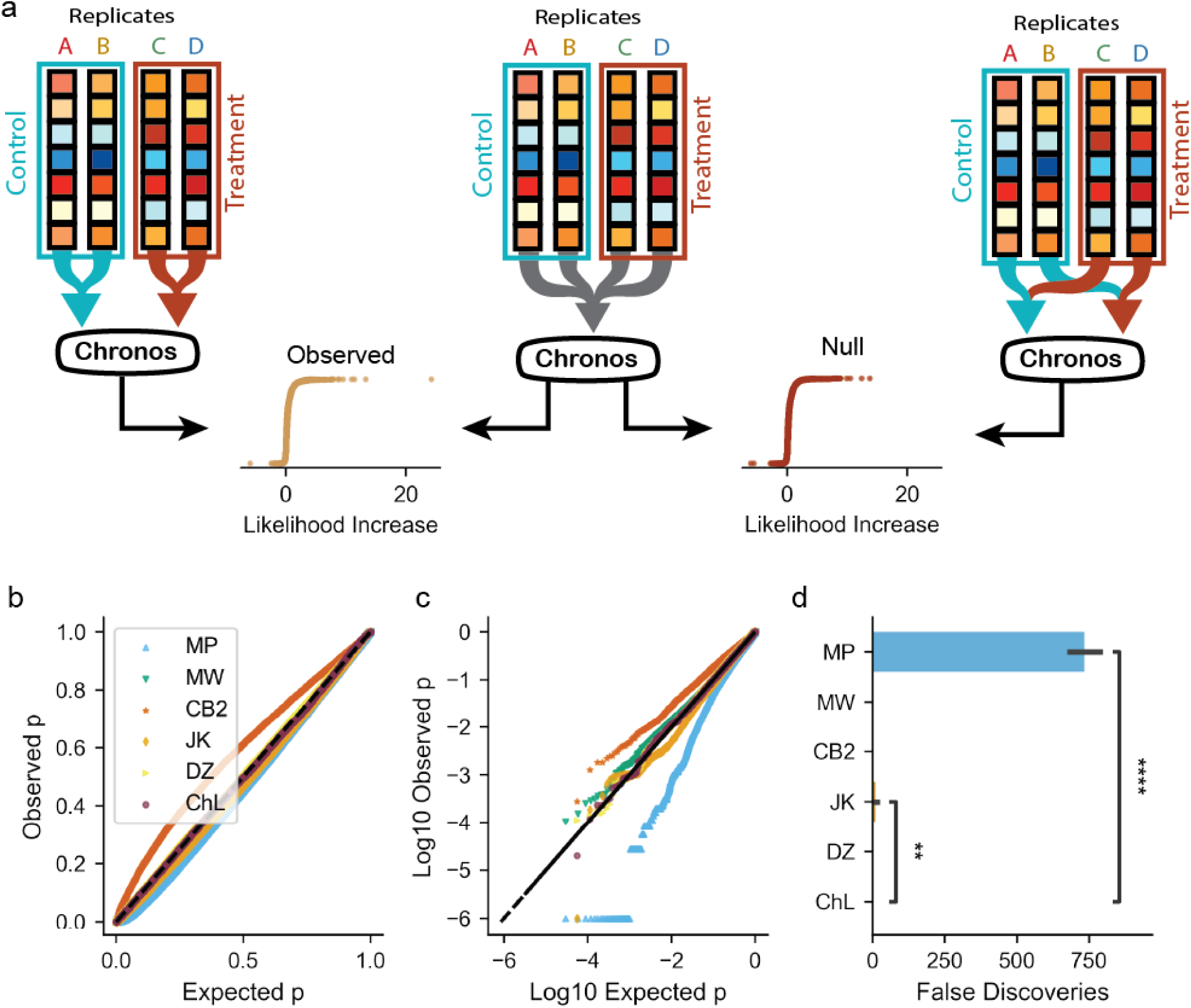
Comparing conditions. **a**. Schematic of the Chronos difference method for constructing a statistic for the alternative hypothesis that the true gene effect is different in two conditions. A model is trained with all replicates of a cell line treated as a single cell line, and the likelihood of the data estimated. Then, a model with the two conditions treated as separate lines is trained. The difference in likelihood between this model and the previous is the observed statistic. Finally, the assignments of replicates to the two conditions are permuted so that the null hypothesis is true, then new models are trained. The difference in likelihood between this model and the undistinguished model is the null distribution. **bc.** *p* calibration. The four replicates in the Avana screen of PSN1 were divided into two pseudo-conditions, so no true differences exist and *p* is expected to be uniform. DZ: DrugZ. **d.** Number of false discoveries with each method. The number of asterisks indicate the significance to that power of 10 using a *t-*test on bootstrap resampling.

In the asymptotic limit, this likelihood statistic should be *χ*^2^-distributed under the null hypothesis(21), but in our tests we found that real data produces likelihoods with longer tails than predicted by *χ*^2^ (**Suppl. Fig. 2a**). Therefore, we must construct an empirical null distribution. To generate data with no true differences between conditions, we exploit the fact that most experiments include at least two replicates for each condition. By swapping replicates between conditions, we create pseudo-conditions that have equal numbers of replicates from the two conditions of interest. As the pseudo-conditions have no genuine biological differences, if we train a Chronos model that distinguishes the pseudo-conditions from each other, the resulting log-likelihood increases can be used as the null hypothesis. For increased power, we train as many models as there exist meaningfully unique permutations (two, if there are only two replicates in each condition). We use some additional approaches to improve power; see Methods.

**Supplementary. Fig. 2:**
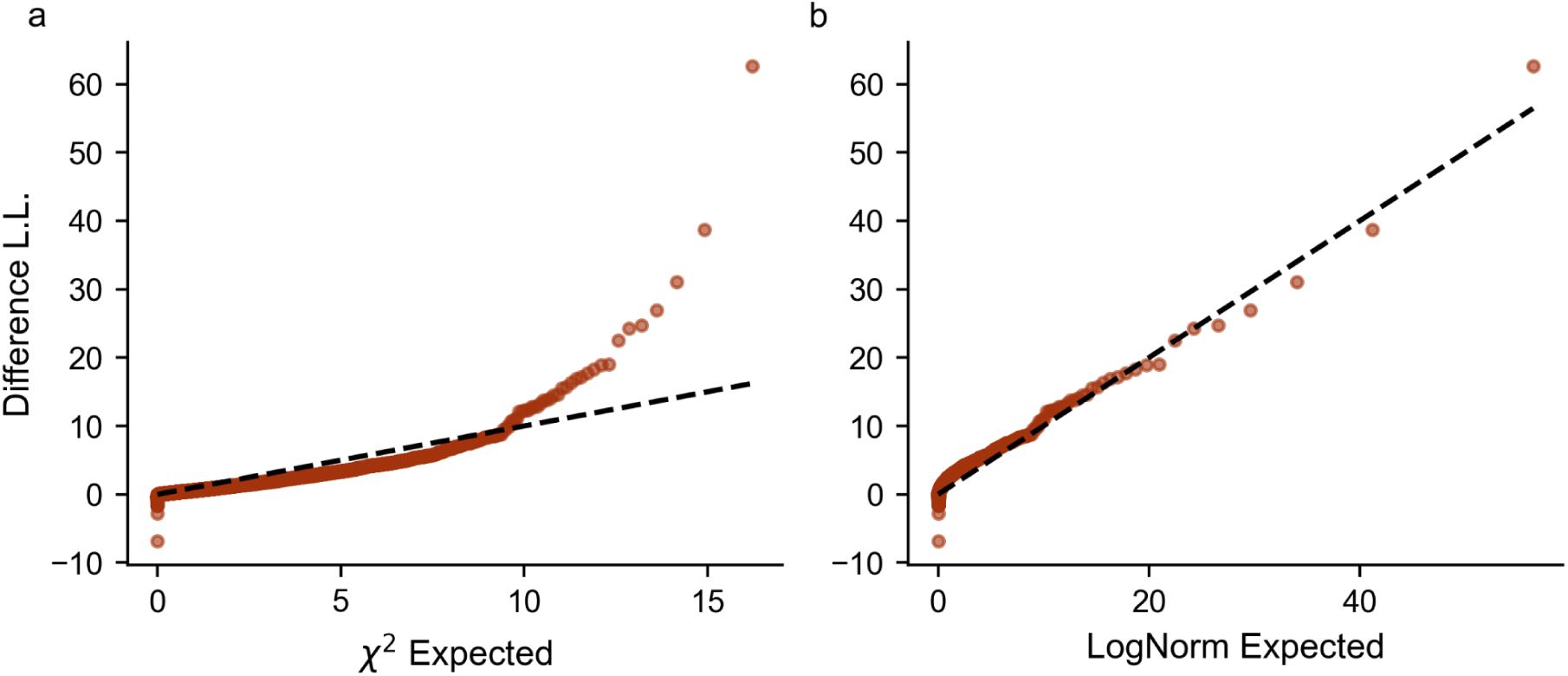
The null distribution for likelihood comparison between condition-distinguished and condition-nondistinguished models. **a**. A quantile-quantile plot of the null distribution found by taking random pairs of replicates of the Avana screen of PSN1 as pseudo-conditions and calculating the increase in log likelihood per gene seen when assigning each pseudo-condition a separate gene effect profile in Chronos, *vs* the *χ*^2^ distribution. **b.** Same for the log-normal distribution.

For these comparisons we added DrugZ, a method that was designed specifically to compare CRISPR screens in two different conditions(14). As before, we start by evaluating each method’s control of false discovery in a setting where there are no true discoveries. The DepMap Avana screen of the PSN1 cell line, which has four biological replicates in the same culture conditions, provides such a setting. We labeled replicates A and B as the treatment condition and replicates C and D as the control condition. With no biological difference between replicates, methods with good control should report no discoveries. We find that Chronos and DrugZ exhibit good calibration of *p*-values and control false discovery in this setting (**Fig. 4b-d**). MAGeCK-Wald also appears to control false discovery, despite performing poorly on the previous task. CB2 *p*-values, which before were optimistic, are now pessimistic, i.e. larger than expected; it reports no false discoveries.

Because the PSN1 test consists of replicates of the same screen, it lacks a realistic representation of the biases that occur when comparing different screens. We took the anchor screens from DeWeirdt *et al*. as a realistic test case(22). These data consist of three cell lines screened in the presence or absence of some perturbation, namely chemical inhibitors of BCL2L1, MCL1, and PARP, or a single *S. aureus* guide targeting the same. Evaluating unexpressed genes, we find a similar pattern of false discovery control among unexpressed genes as in the PSN1 case, with Chronos, DrugZ, and MAGeCK-Wald appearing well-calibrated, JACKS and MAGeCK-permutation being optimistic, and CB2 being pessimistic (**Suppl. Fig. 3**). However, MAGeCK-Wald and DrugZ reported many more discoveries than Chronos despite their improved calibration. We examined these purported discoveries in the OVCAR olaparib condition and found that the MAGeCK and DrugZ discoveries, unlike Chronos, are heavily skewed towards the strongest common essential genes (**Fig. 5a**), and in fact MAGeCK-Wald reports the majority of all strong common essentials as significantly different between olaparib and control **(Fig. 5b**). As these gene knockouts are rapidly cell lethal in all conditions, it is unlikely that they have a meaningfully increased viability effect in olaparib vs the control condition. Rather, the olaparib arm appears to have achieved more sensitive detection of dropout than the control condition (**Fig. 5c**). This is one of the types of screen quality artifacts often seen in CRISPR experiments(15), so we interpret these highly lethal knockouts as uncontrolled false positives. We took the 30 gene knockouts with median Chronos gene effect scores ≦ -2 across all the DeWeirdt screens as universally lethal, and examined the calibration of each method on this set. Only CB2 and Chronos (0 false discoveries) controlled false discovery on this set within the DeWeirdt condition comparisons (**Fig. 5 def**).

**Figure 5:**
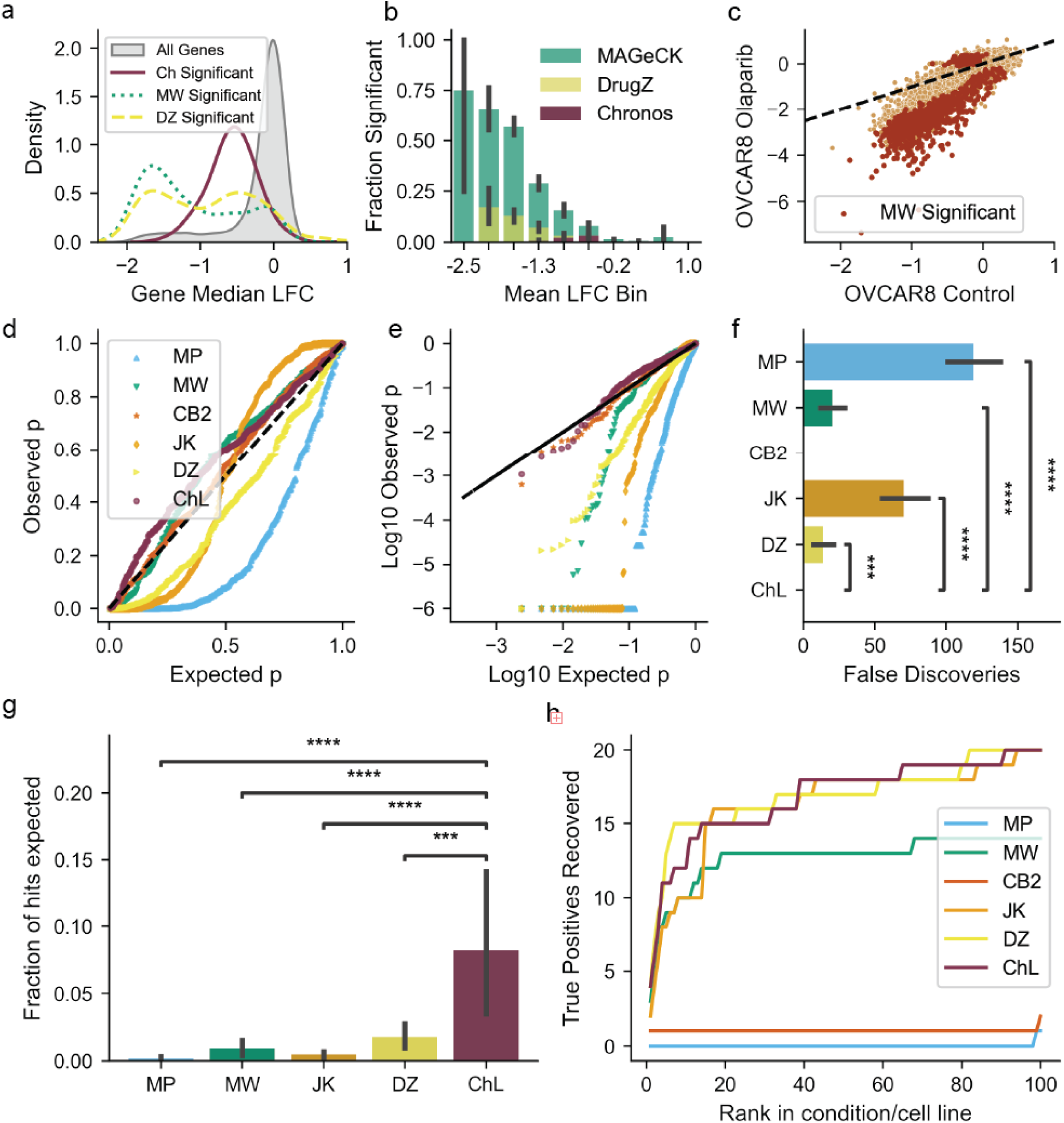
DeWeirdt isogenic and drug screens. **a.** Distribution of mean log fold-change (LFC) in gene abundance for all genes, those found differential by Chronos, and those found by MAGeCK-Wald and DrugZ. A gene scoring in multiple conditions is included multiple times. **b**. Genes are binned by mean LFC over the whole dataset, and the fraction found significantly different in OVCAR8 Olaparib by MAGeCK-Wald, DrugZ and Chronos reported for each bin. **c.** Comparison of the OVCAR8 LFC for each gene in Olaparib vs the control condition. **de**: *p*-value distribution for universally lethal genes to have differential gene effect in the three lines screened by DeWeirdt *et al*. across all conditions and cell lines vs control (*n* = 420). **g**: For all results reported as significant at FDR < 0.1, what fraction are expected hits. The number of asterisks indicate the significance to that power of 10 using a *t-*test on bootstrap resampling, while error bars show 95% CIs. **h.** The number of expected hits recovered in *N* most significant results for each method, for *N* ∈ [1, 100].

**Supplementary. Fig. 3:**
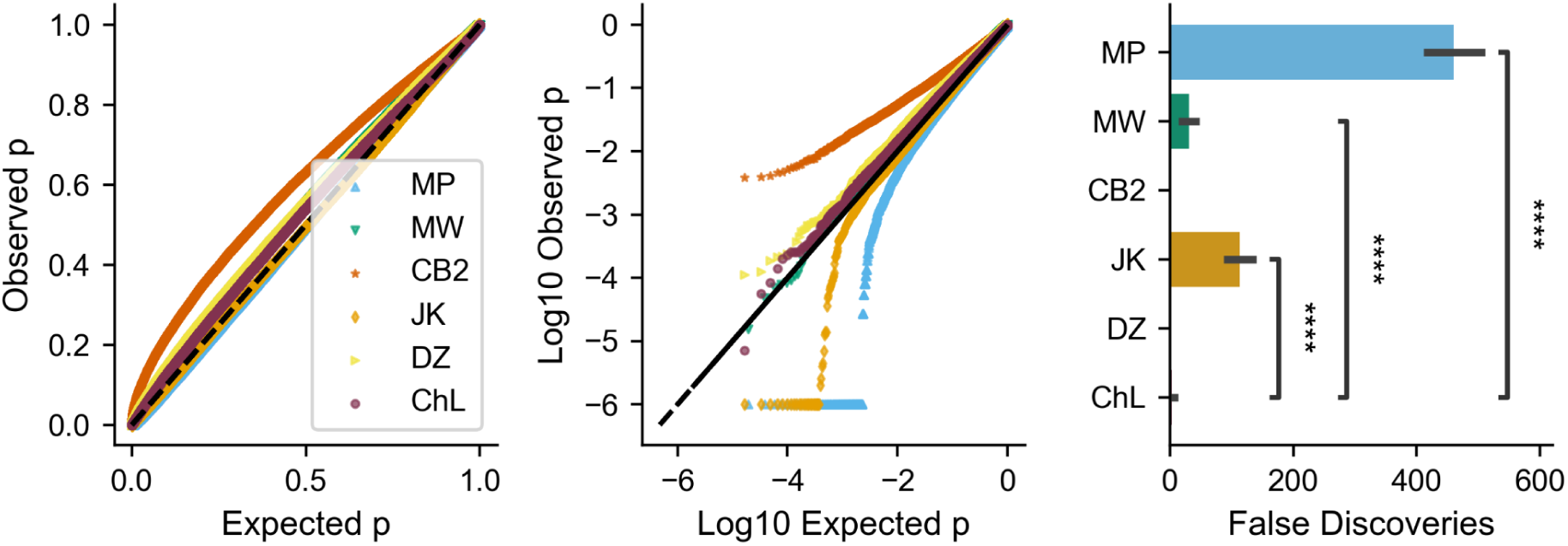
Calibration on negative controls in DeWeirdt data. *p*-value calibration and false discoveries in the DeWeirdt condition comparisons using the test null distribution across all combinations of cell line, condition, and unexpressed genes (*n* = 61,393). Total false discoveries at FDR < 0.1 are reported for each method. The number of asterisks indicate the significance to that power of 10 using a *t-*test on bootstrap resampling.

As the DeWeirdt dataset includes perturbations with known synthetic lethal partners, we can also evaluate each method on its ability to identify a limited set of true positives. Joint inhibition or loss of MCL1 and BCL2L1(23,24), BCL2L1 and MARCH5, UBE2K, or UBE2J2(25), BCL2L1 and BAX(26), or PARP and BRCA1/2(27), are established vulnerabilities in cancer. We evaluated the recovery of these pairs in the DeWeirdt data using both the chemical and *S. Aureus* perturbations in each cell line for 28 true positive combinations in total (**Suppl. Fig. 4**). Unsurprisingly, the rank of algorithms by the total number of true discoveries largely follows the rank by the number of false discoveries or total claimed discoveries; MAGeCK-permutation reports 8,963 discoveries and recovers 20/28 true positive combinations, while CB2 reports no discoveries of any kind. The exception is DrugZ, which reports fewer total discoveries than MAGeCK-Wald (837 vs. 975) but more known discoveries (15 vs 8). For a researcher, an algorithm that recovers almost all true hits but only among a list of thousands of putative hits obscures rather than reveals the key biology. Therefore, we examined how efficiently different methods enriched for known synthetic lethals among reported hits. Among hits at 10%FDR, Chronos was most efficient at 7 expected hits of 78 reported hits, followed by DrugZ at 15 expected hits among 837 findings (**Fig. 5g**). If one ranks all results by FDR and takes the top *N* results for followup, DrugZ, Chronos, and JACKS all recover similar numbers of real results up to *N*=100, while MAGeCK-WALD has noticeably worse performance and MAGeCK-MP and CB2 recover almost no known hits in the top 100 (**Fig. 5h**). It is interesting to note that, while most methods identified BAX as significant hit with BCL2L1 inhibition in both cell lines, in all cases the direction of the effect was the opposite of what we expected from literature: BAX knockout appeared to be a resistance mechanism in these screens, rather than sensitizer as reported by Lopez *et al*.(26). These results were counted as misses when scoring the algorithms.

**Supplementary. Fig. 4:**
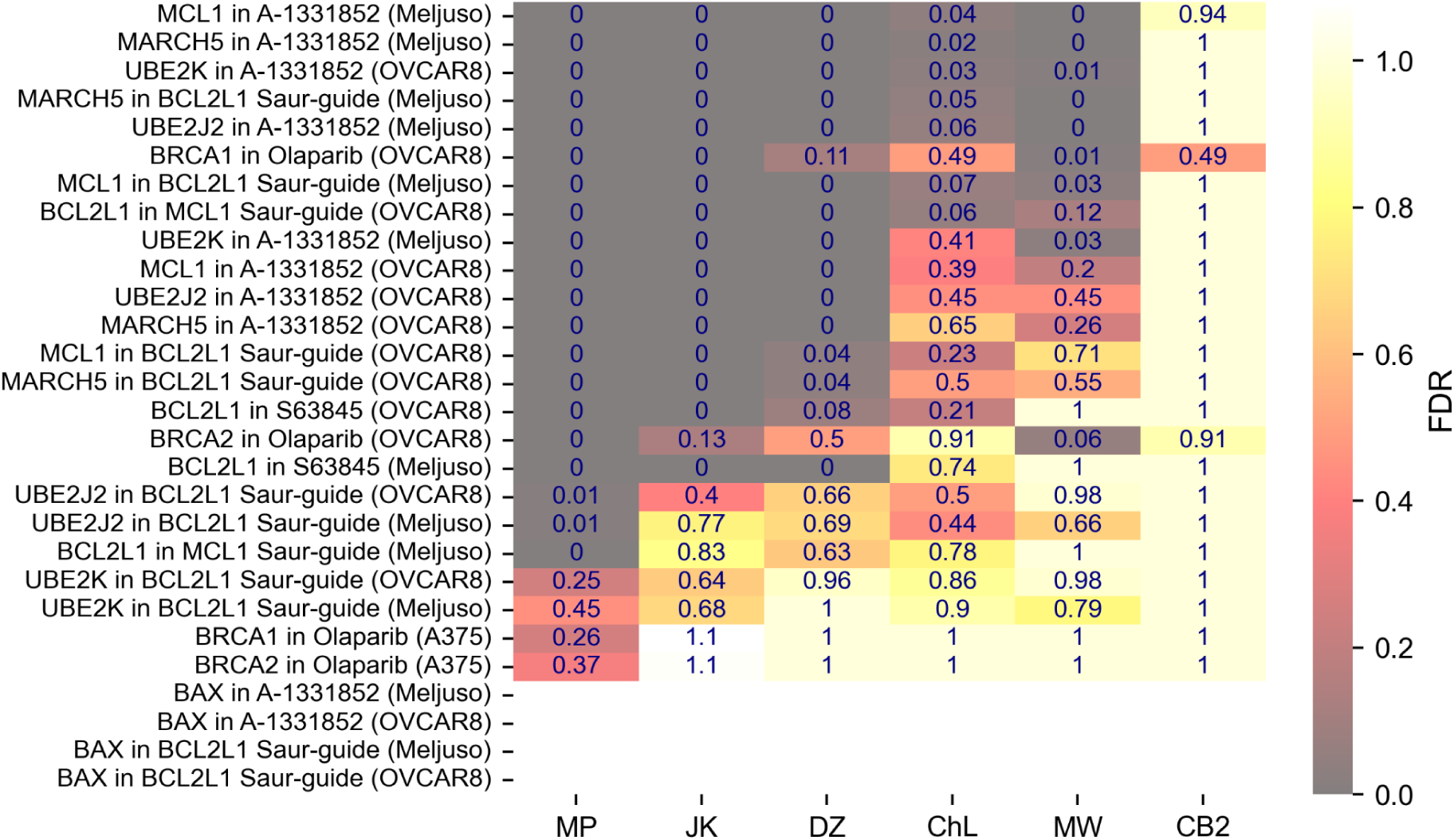
Expected synthetic lethal hits in the DeWeirdt screens. For a curated list of treatment/knockout pairs with expected synergistic interactions, the FDR reported by each method. Empty cells indicate that the effect direction is the opposite of expected (antisynergistic rather than synergistic), so the reported FDR has been NAed.

## Discussion

In this paper, we focus on two tasks: identifying genes whose CRISPR knockout causes a true loss of viability; and identifying genes whose knockout has differential effects on viability between two conditions. We introduced methods that work in concert with the Chronos algorithm to make significant finds for both tasks, and tested their performance against existing methods. On the first task, we find all existing methods fail to correctly control false discovery as they systematically underestimate the left tail of the null distribution, while both the frequentist and empirical-Bayesian Chronos methods succeed. If good positive control genes can be identified, we recommend the empirical-Bayesian method as it is better calibrated and exhibits more power with fewer controls needed. On the second task, we find that MAGeCK-Wald and DrugZ are able to control false discovery except among common essential genes. Their success is notable, as both methods make a similar assumption: that appropriately normalized log fold changes (DrugZ) or the closely related *β* scores (MAGeCK-Wald) are normally distributed under the null hypothesis. Since the difference between the same knockout in two conditions is generally smaller than the difference between pDNA and a late timepoint, it may be that this assumption holds better in task 2 than task 1. However, screens in two different conditions will often have different screen quality, which causes both MAGeCK-Wald and DrugZ to systematically report universally lethal genes as differential between conditions. In contrast, Chronos’ likelihood test controls false discovery even in the presence of screen quality bias. MAGeck-Wald also performs notably worse than Chronos and DrugZ at surfacing known hits near the top of its rankings.

However, Chronos’ methods have important limitations. Both methods for identifying dependencies in single conditions require identifiable gene controls–either a very large set (preferably > 500) of negative controls for genome-wide screens with the frequentist method, or high confidence sets of negative and positive controls for the Bayesian method. For identifying differential viability between conditions, Chronos requires at least two independent biological replicates of each cell line in each condition. More subtly, Chronos depends on the two replicates being genuinely independent. If the replicates are coinfected before splitting, this assumption is violated, and Chronos’ estimated *p*-values are no longer guaranteed. We have included a check for whether biological replicates appear not to be independent which will generate a warning for the user (see Methods).

Supposing an experiment does not meet the criteria for the Chronos methods introduced here, what should researchers use to assess significance? For the first task, we suggest that, if the experiment lacks adequate controls to use either of Chronos’ methods, researchers should not attempt to report a significance and instead focus on independently validating the strongest findings. However, for the second task, we believe DrugZ is a good choice, provided that the user either verifies that there is no screen quality bias or is willing to exclude common essential genes before running the algorithm. We recommend it over MAGeCK-Wald, since it has better control of false discovery and better sensitivity. It is also much faster to run.

Given the widespread use of MAGeCK to analyze CRISPR screens(3) and the miscalibration of its *p*-values as reported here, one may reasonably worry how contaminated the literature is with false positives. We believe the problem is not as severe as it might seem. Researchers have adopted a number of heuristic strategies which reduce false positives. These include filtering for hits that appear logical (or exciting) based on existing biological knowledge, taking the intersection of multiple screens, and limiting consideration to the results with the strongest effect sizes. This means that the majority of the “significant” findings nominally discovered in a screen are not pursued in follow-up experiments. However, these ad hoc filters limit a study’s power for true discovery and introduce an uncomfortable amount of leeway for scientists to cherry-pick favorable results. Although no method can substitute for scientific judgment, we hope that introducing more rigorous control of false discovery will increase researcher confidence in reportedly significant findings. This in turn can empower scientists to pursue more novel or unexpected findings in their experiments.

## Conclusions

Existing methods for analyzing CRISPR have not been proven to control false discovery. In this study, we introduced a new python submodule, Chronos.hit_calling, with a suite of tools for assessing significance in CRISPR screens. We evaluated these tools together with existing methods and found that only Chronos’ *p*-values are consistently calibrated. Chronos’ estimates of false discovery risk are more accurate than competitors and better separate true from false positives. When tasked with identifying differential viability impacts between conditions, we found that prior methods tended to report many highly lethal knockouts as differential between conditions despite a lack of biological plausibility. Chronos avoids this problem while still capturing true differential viability. The code for Chronos and a jupyter vignette illustrating its use can be found on Github(28).

## Methods

### Controlling false discovery of dependencies

Chronos’ hit_calling package provides two methods for estimating false discovery of genes with negative viability effects on knockout. The frequentist method uses only negative control genes. For any given query gene, we count the *n* negative controls with more negative gene effect scores than the gene in question, out of *N* total negative controls. The *p*-value is then simply (*n*+1) / (*N* + 1), and the FDR is estimated by the two-stage Benjamini-Hochberg procedure(17).

In the Bayesian method, we first construct density estimates of the positive and negative controls gene effect distributions. The kernel width is chosen by Scott’s method(29). These densities are then projected onto a grid extending past the most extreme observed gene effect scores in both tails. Because the density of a gaussian kernel density estimator can fall to 0, we force the negative control distribution to have a minimum density of 10^-16^ in the right tail, and the positive control distribution to have a minimum density of 10^-16^ in the left tail. The thresholds for the tails are -3 and 1 by default. The grid of values is then interpolated by scipy’s interpolate function.

We then set the mixing ratio of genes belonging to the negative (*n*) or positive (*p*) control distribution, by default 0.5 for each; i.e., we start with an equal prior that genes are either dependencies or not. The posterior probability for an observation of gene *g* with gene effect *G*_*g*_ to be drawn from the positive distribution is:

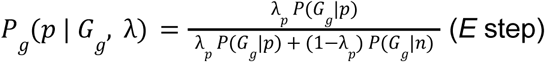

After computing all *P*_*g*_, we then update the estimated fraction of genes estimated to be hits:

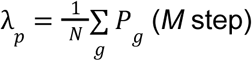

This is repeated until convergence, meaning the fractional increase in the likelihood of the data is less than some specified value (10^-7^ by default). The resulting probabilities can still show odd tail behavior due to finite sample size. In particular, because there may be fewer positive controls than negative controls and the positive control distribution is wider, the kernel size for the density estimator for positive controls is often significantly wider than for the negative controls. This means that points in the right tail of the distribution can have an increasing probability of being generated by the positive control distribution. Because the probability of dependency must decrease monotonically with increasing viability, we fit a generalized linear model (GLM) using the binomial distribution and logit linker function to the empirically estimated probabilities, excluding points more negative than the most negative positive control and more positive than the median of negative controls. The predictions of the GLM show a close match in the transitional region between the positive and negative controls, while eliminating the low density artifacts in the tails (**Suppl. Fig. 5**), and are used as the probability. The estimated number of true discoveries among all genes with gene effect below some threshold is the sum of their probabilities of dependency. Accordingly, we can estimate the positive false discovery rate for some gene as 1 - the sum of the probabilities for all genes with equal or more negative gene effect(18).

**Supplementary. Fig. 5:**
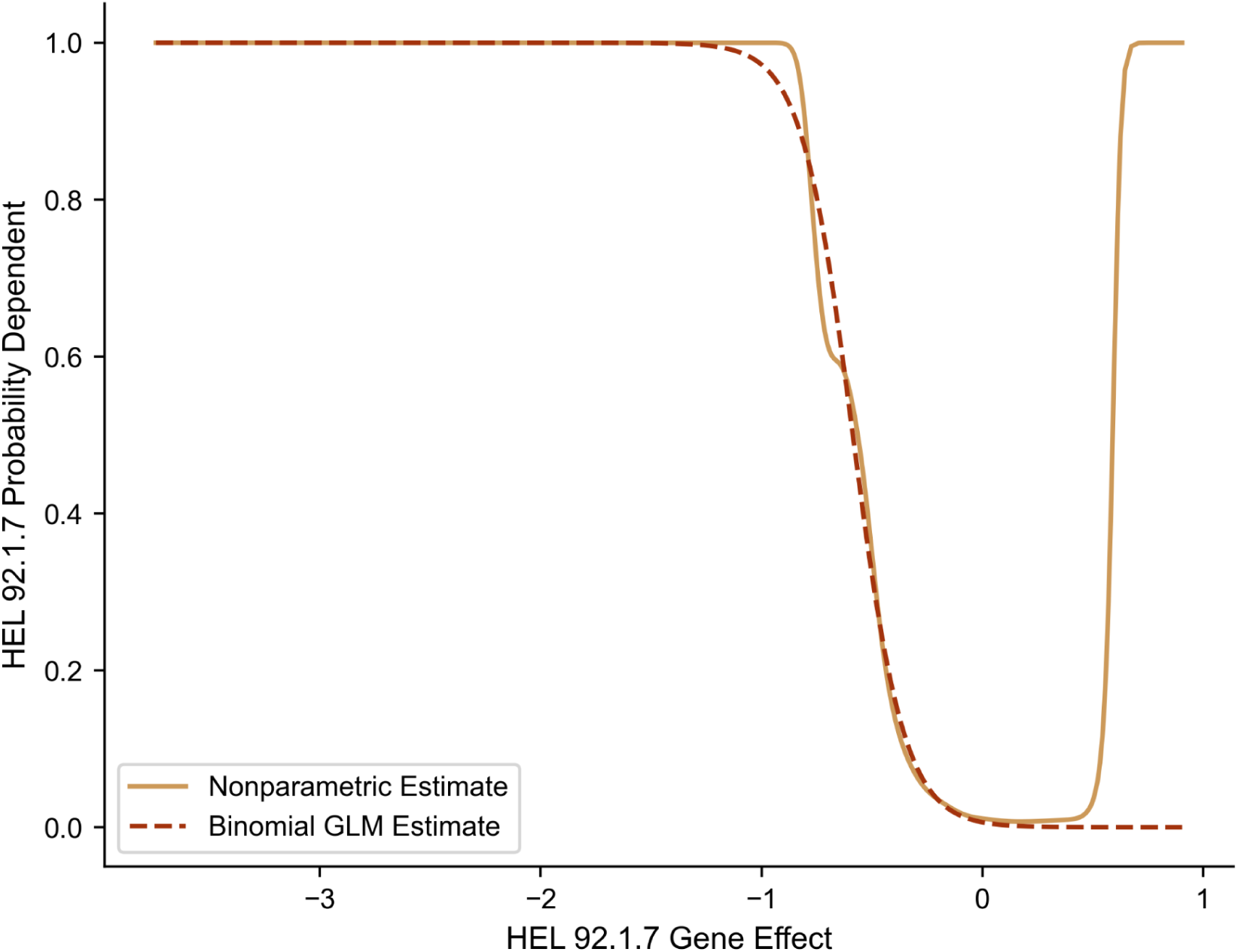
Example of the effect of applying the binomial generalized linear model to the nonparametric estimate of dependency probability.

### Controlling False Discovery of Differential Viability

Suppose we have a cell line screened in two conditions, which for convenience we’ll call treatment and control. We observe different gene effects in each condition. We wish to choose the largest possible set of gene knockouts such that no more than *X* % have no true difference in viability effect between the conditions. For this task, we take as our statistic the change in the log likelihood of the data (according to Chronos) between a model where all conditions are constrained to have the same gene effect (the undistinguished model) and a model where each cell line in each condition is permitted to have a distinct gene effect (the distinguished model).

Specifically, for each gene and cell line, we sum the log likelihood of all sgRNAs targeting the gene among all replicates for the cell line in both conditions according to a model that treats the two conditions as separate cell lines, *vs* one that treats them as a single cell line. To ensure that only differences in gene effect can contribute to an increase in likelihood, all parameters including the mean effect of each gene knockout are learned using the undistinguished model. For distinguished models, only the departures of the gene effect from the mean are relearned. Under the null hypothesis, this statistic should asymptotically approach a *χ*^2^ distribution with one degree of freedom(21), but in a strictly null test case we find a notably longer positive tail (**Suppl. Fig. 2a**). This may be due in part to the fact that different gene effects are coupled by multiple mechanisms in the Chronos model. To capture the long tail, we need to generate an empirical null distribution. If we create two pseudo-conditions, each with an equal number of replicates (of each cell line) drawn from the two real conditions of interest, there are no true differences in gene effect between the pseudo-conditions. Therefore, we take the distribution of change in likelihood of the distinguished over the undistinguished models using these pseudo-conditions as the null distribution.

To create the pseudo-conditions, for each cell line we take *n* replicates from each condition, where *n* is the largest integer ≤ the smallest number of replicates available in either condition over all cell lines. For example, given 3 replicates in the treatment condition and 4 in the control, we can only use 2 replicates from each and still construct 2 pseudo-conditions with equal numbers of replicates from replicate and control. We then generate all possible single partitions of these replicates that have equal numbers of treatment and control replicates on each side of the partition. The two halves of the partition are the pseudo-conditions. To avoid redundant sampling, we exclude partitions that are mirror images of an existing partition. For example, if we have a partition

Treatment replicate **A**, Control replicate **A** | Treatment replicate **B**, Control replicate **B**

we exclude the partition

Treatment replicate **B**, Control replicate **B** | Treatment replicate **A**, Control replicate **A**

from our analysis. The partition is performed *per biological replicate*, not per sequence. Thus, if replicate A has three late timepoints that were sequenced, all three timepoints will always be on the same side of the partition.

For each remaining pair of pseudo-conditions, we compute the same increase in log-likelihood statistic. We then pool statistics across the different partitions and genes to increase statistical power. The empirical *p*-value for any gene to have a difference in gene effect between conditions in a cell line is the number of statistics from the pseudo-conditions with equal or greater increase in likelihood + 1, divided by the total number of such statistics + 1.

Like any empirical *p*-value, the achievable significance is strictly limited by the number of samples in the null. Ideally, we would like points that are more extreme than the most extreme null value to have even smaller *p*-values, which requires some parameterized model for the empirical null distribution. We have found that we can fit a log-normal distribution to the right tail of the null with good fidelity **(Suppl. Figure 2b**). We want the fit to be best for the extreme right tail, as that is where we will rely on it for extended *p*-values. We use a simple linear regression on the 20 gene-cell line pairs with the largest increases in likelihood from the null distribution. The endogenous variable *y* is the log increase in likelihood for the pairs:

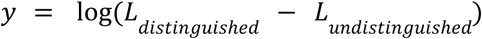

The exogenous variable *x* is the expected value for those pairs according to a unit normal distribution:

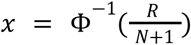

where Φ^−1^ is the inverse cumulative density function of the unit normal distribution, *R* is the rank of the tested pair among all pairs, and *N* is the total number of such pairs. In the regression

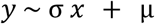

we weight each sample by its increase in log likelihood. We then assume that the log of the null distribution is the Gaussian *N*(µ, σ). For likelihoods in the true condition that exceed the largest likelihood found in the null distribution, we substitute the *p* value found from this log-normal approximation if it is smaller than the empirical *p*-value.

Gene effect estimates for genes with very low total readcounts (summed over guides) in the initial infection population are intrinsically noisier than the majority of genes. To prevent these genes from being assigned inflated *p*-values, and to keep their high variance from creating pessimistic *p*-values for the remaining genes, we segregate genes in the bottom 5% of total reads in the infection pool from the remaining genes before computing *p*-values.

Once *p*-values have been computed, FDRs are computed from them with the two-stage Benjamini-Hochberg procedure(17). All *p*-values and FDRs are computed separately per cell line if multiple cell lines are present.

If replicates are not biologically independent, this method can fail to control false discovery. For example, if the replicates in a given condition share a bottleneck, the genes affected by the bottleneck will look systematically more or less depleted in the condition, perfectly confounding the biological effects of the condition. This can be detected using negative controls which are expected to have no true differential signal. If the negative controls for replicates within conditions resemble each other much more than between conditions, this indicates confounding. Chronos checks the correlation matrix of replicates using only negative controls and warns users if mean within-condition replicate Pearson correlation is at least 0.1 higher than the cross-condition correlation for any condition.

If there is confounding, Chronos will still attempt to ensure its *p*-values are calibrated. Given a set of negative control genes, Chronos will check that *p*-values for those genes follow the expected uniform distribution using a KS test. If the KS statistic is greater than

### Other changes to the Chronos model

Numerous changes have been made to Chronos since the original publication(15). These changes improved Chronos’ performance in settings with high overdispersion or multiple libraries and introduced the ability for Chronos to load parameters learned from previous runs and use them to process new screens, as well as plotting and QC report modules and improved data normalization. Some additional changes introduced with this work are noted below.

Chronos estimates two screen-quality related terms: the unperturbed relative growth rate *R*_*c*_ and the maximum penetrance of a CRISPR knockout genotype *p*_*c*_. The *c* in each term indexes the cell line, since these terms previously were estimated per cell line. However, biological replicates of the same cell line screened in different conditions can have very different values for these parameters. Accordingly, we now estimate these parameters per biological replicate, which is never present in more than one condition. If the user does not specify the biological replicate in the sequence map, Chronos assumes every late timepoint represents a distinct replicate.

### Evaluation and Analysis

Notebooks containing the analyses performed in this manuscript are provided in the supplementary materials.

Data for the Avana screens was downloaded from the DepMap 23Q4 data release(30). Data for the DeWeirdt screens was taken from DeWeirdt *et al.*(*22*).

To isolate the direct effects of the algorithms, all methods were presented with identically normalized data. All guides targeting multiple genes were excluded. Data was normalized by running `chronos.normalize_readcounts`, followed by `chronos.nan_outgrowths`. For Avana screens, these were run on the entire set of DepMap screens before subsetting. After subsetting, genes with any null values were dropped. No copy number correction was performed either before or after running. Training positive and negative controls were taken from AchillesCommonEssentialControls and AchillesNonessentialControls from that dataset, respectively(30,31). For a test set of negative controls, we used genes that were unexpressed in all cell lines examined, then excluded the training set of negative controls. For a test set of positive controls, we chose genes whose 90th percentile most positive gene effects were below -0.2 in both KY(32) (315 Cas9 screens integrated into DepMap) and Humagne(33) (15 AsCas12a screens integrated into DepMap)(30), excluding genes in either the negative or positive control training sets. Using transcript per million (TPM) data in 23Q4, protein coding genes were considered unexpressed if log2(TPM+1) < 0.1. For all methods, we used the false discovery rate estimates provided by the method. Throughout, we have considered FDR < 0.1 the threshold to reject the null hypothesis.

To evaluate MAGeCK, MAGeCK-MLE 0.5.9.5 was installed and run on data with commands of the form:

~~~
mageck mle -n {directory_location} -k {counts_file_location} -d
{design_matrix_location} --control-gene
{negative_control_gene_file_location} --max-sgrnapergene-permutation 5
--permutation-round 4 --threads 2
~~~

The input and output files from MAGeCK commands are provided in.

To run CB2, we used version 1.3.4 with the functions `measure_sgrna_stats` and `measure_gene_stats` with default parameters.

To run JACKS, we first downloaded version 0.2 from GitHub(34). Certain functions that the JACKS module imported from scipy were subsequently moved to numpy; we edited JACKS to import from the correct package. Additionally, in its existing form, JACKS computed FDRs but did not have a parameter to report the actual value, so far as we could determine; the code was edited to report the values computed. The edited versions of `jacks_io.py` and `infer.py` are provided in Figshare(35). JACKS was trained by first saving readcounts, sequence maps, guide maps, and control genes in the expected format, then calling `jacks_io.runJACKS` with the specified files and `fdr=1.01, n_pseudo=2000`.

To run DrugZ, we downloaded commit eb15d34 from Github(36). The code used to run it in python is provided in Figshare(35). We used default parameters.

Chronos methods `chronos.Chronos(…).train` and

`chronos.hit_calling.ConditionComparison(…).train` were run with the default parameters in all cases.

The 5 cell lines chosen for the Avana evaluation were ACH-000077: MJ, ACH-000093: Panc 05.04, ACH-000101: KE-37, ACH-000124: OCI-LY-19, ACH-000147: T-47D. These were chosen to be screens with expression data present, then the lowest DepMap model ID. To evaluate FDR control in a setting with no true positives, we subsetted for genes in the negative control list or unexpressed in all five lines, then ran the considered methods. The negative control list was used as the training set of negative controls, which were passed to those methods that accepted negative controls, while the unexpressed list (excluding the training list) were used for evaluation. For MAGeCK and CB2, we provided design matrices to compare final sgRNA abundance to pDNA. We plotted the *p*-value distribution for gene-cell-line pairs from each method against the expected (uniform) distribution, and computed the total false discoveries (all genes found to have FDR < 0.1). With no positive controls available, the Chronos Bayesian method could not be used.

We then trained each method with the same five lines but all genes included. We used the same partitioning of negative controls, and randomly split the positive controls into equal test and train groups. As the discoveries now include true discoveries, we reported the fraction of discoveries at FDR < 0.1 known to be false (using expression, excluding the training negative controls). Using the test sets of positive controls and the unexpressed genes in each cell line, we computed precision and recall for each method using possible thresholds on its effect size estimate (gene effect for Chronos, *β*-scores for MAGeCK, and log fold change for CB2) as the statistic for calling discoveries.

To assess FDR calibration, we trained with the five Avana screens once more, now including only positive and negative controls. Both control groups were split evenly into train and test sets. For each method, gene and cell line pairs were ranked in order of increasing FDR estimate and the true FDR for each gene computed as the fraction of all genes with equal or lesser FDR that were negative controls, only the test control genes.

To evaluate false discovery control for differences between viability in two conditions, we first took the PSN1 Avana screen, which has 4 replicates. We assigned the first 2 to be a treatment condition, and the second 2 to control. We then compared these two “conditions” with each method. We examined *p*-value calibration and false discovery control as before, again with the assumption that no true viability differences should exist. As Chronos does not use control groups to assess significance for this task, no train-test split was performed.

We then evaluated the DeWeird isogenic screens by training each available alternate condition (either a drug treatment or an *S. aureus* guide) against the untreated condition for the cell line. We tested false discovery among negative controls as before. We then binned genes into 10 bins of equal width according to their median log fold change across all the cell lines and conditions in this dataset. Within each bin, we examined the percentage reported significantly different in the OVCAR cell line in olaparib vs control by either MAGeCK-Wald or Chronos. To test the recovery of known hits, we took the well-established synthetic lethal pairs: BCL2L1/MCL1, and PARP/BRCA1 or 2. We considered a genetic knockout of one half of the pair to be an expected hit if the other half was either chemically inhibited or knocked out with *S. aureus* cas9. The exception is BRCA1/2 knockout with PARP1 knockout. Knocking out PARP1 alone is not phenotypically equivalent to the dual PARP1/PARP2 inhibitor olaparib.

## Code availability

The Chronos methods used in this manuscript are available at Github(28). Notebooks used for running methods and analysis are available in the supplementary information and in Figshare(35).

## Declarations

### Ethics Approval and Consent to Participate

Not applicable

### Consent for Publication

Not applicable

### Availability of Data and Materials

The datasets supporting the conclusions of this article are available in the Figshare repository(35).

### Authors’ Contributions

J.D. designed the study, developed the code, and drafted the manuscript. A.K proposed the likelihood test for task 2 and edited the manuscript. B.d.K. wrote and corrected code and edited the manuscript. F.V. and C.D.C. supervised the project and edited the manuscript.

